# MyoAssist 1.0: An Open-Source Framework for Neuromechanical Simulation of Physical Human-Device Interaction

**DOI:** 10.64898/2026.08.25.746839

**Authors:** Calder Robbins, Hyoungseo Son, Chun Kwang Tan, Cheryl Wang, Roger van Kanten, Massimo Sartori, Guillaume Durandau, Vikash Kumar, Vittorio Caggiano, Seungmoon Song

**Author notes:** Co-first authors (Calder Robbins, Hyoungseo Son). Corresponding author: S. Song.

## Abstract

Physical human-device interaction is central to many emerging technologies in neurorehabilitation and assistive robotics, but simulation-based research in this area remains fragmented across musculoskeletal models, assistive-device representations, task definitions, and controller-development workflows. This fragmentation limits the accessibility, reproducibility, and extensibility of studies on prostheses, exoskeletons, wearable rehabilitation devices, and related human-device systems. Here we introduce MyoAssist 1.0 [1], an open-source framework for neuromechanical simulation of physical human-device interaction built within the MyoSuite ecosystem. MyoAssist organizes each simulation environment as a composed human-device-task system that combines compatible musculoskeletal, assistive-device, and task-scenario components through a shared composition pipeline. The current release includes 15 assistive-device models spanning gait assistance, upper-body support, manipulation, and seated mobility and supports compatible musculoskeletal models ranging from reduced lower-limb models to a 416-muscle full-body model. These human-device systems can be simulated within the broad task scenarios provided by MyoSuite, while MyoAssist adds locomotion-specific task scenarios with configurable terrain and target-velocity conditions for gait-assistive studies. MyoAssist also provides two complementary controller-development frameworks: a reinforcement-learning framework for training adaptive policies and a controller-optimization framework for tuning structured, interpretable human and device controllers. Both frameworks operate on the same simulation environments and provide standardized evaluation outputs for inspecting, comparing, reusing, and extending learned and structured control strategies. By integrating modular human models, assistive-device models, task scenarios, and training workflows under a shared open-source interface, MyoAssist aims to lower the barrier to reproducible simulation-based research and to support collaborative development of assistive technologies for neurorehabilitation and physical human-device interaction.

## I. Introduction

Physical human-device interaction lies at the core of many emerging technologies in neurorehabilitation and assistive robotics, including powered prostheses, exoskeletons, wearable rehabilitation devices, and intelligent mobility systems. These systems operate through tightly coupled interactions between neural control, musculoskeletal dynamics, assistive-device mechanics, and environmental conditions, requiring coordinated adaptation across both biological and robotic components. Developing and evaluating such systems experimentally is often expensive, time-consuming, and difficult to scale across users, tasks, and device configurations. As a result, computational simulation frameworks capable of reproducing physical human-device interaction are increasingly recognized as important tools for rehabilitation engineering, motor control research, and assistive technology development [2].

Recent advances in computational musculoskeletal and neuromechanical simulation have substantially expanded the ability to investigate movement, motor control, and assistive-device interaction through physics-based simulation [2]. Open-source frameworks such as OpenSim [3], [4], SCONE [5], DART [6], and MyoSuite [7] provide complementary capabilities for musculoskeletal modeling, predictive simulation, trajectory optimization [8], and learning-based neuromechanical control [9]. These tools have expanded the role of simulation-based assistive-device research beyond retrospective biomechanical analysis to also support predictive design and controller synthesis for exoskeletons, prostheses, and other wearable robotic systems [10]–[18]. These advances are positioning simulation as an engineering front end for assistive technology development, enabling candidate devices and controllers to be explored before extensive hardware prototyping or human-subject testing.

Despite recent advances, assistive-technology simulation remains split across musculoskeletal models, device representations, software ecosystems, and control workflows, creating barriers to multidisciplinary research on coupled human-device systems [2]. Biomechanics studies often emphasize musculoskeletal analysis and trajectory optimization, robotics studies often emphasize hardware modeling and control design, and machine-learning studies often emphasize policy optimization in simplified or study-specific environments. As a result, adapting a simulation to a new device or task, reproducing a method in another laboratory, or comparing biomechanical, robotic, and learning-based control approaches can require substantial redevelopment of model components, interfaces, and training pipelines. These barriers limit accessibility, reproducibility, and extensibility, especially for learning-based and simulation-to-hardware studies, where strong conclusions require transparent models, executable code, and standardized evaluation workflows [12], [19]–[21].

To address these challenges, we present MyoAssist 1.0, an open-source framework for neuromechanical simulation of physical human-device interaction. Built within the MyoSuite ecosystem [7], MyoAssist integrates musculoskeletal models, assistive-device models, task scenarios, and training workflows under a shared computational interface. The current release includes 15 assistive-device models across prosthetic, exoskeleton, upper-body support, and seated-mobility applications and supports musculoskeletal models ranging from reduced lower-limb models to a 416-muscle full-body model. MyoAssist further enables modular composition of human, device, and task-scenario components, allowing researchers to interchange assistive devices, vary musculoskeletal model complexity, and retarget controllers across tasks without re-building simulation pipelines. The framework additionally provides complementary reinforcement-learning and controller-optimization workflows within the same environment interface, supporting both adaptive data-driven policy learning and structured physiologically grounded controller design. Through this unified framework, MyoAssist aims to facilitate reproducible and collaborative research on broader physical human-device interaction problems.

## II. Overview of MyoAssist

MyoAssist is a MyoSuite sub-suite for neuromechanical simulation of physical human-device interaction.

### A. MyoSuite ecosystem and MyoAssist 0.1

MyoSuite is an open-source neuromechanical simulation suite built on MuJoCo [22] with Gym-compatible [23] interfaces. It provides musculoskeletal models and task environments for studying muscle-driven movement, contact-rich interaction, and controller development in physics-based simulation. MyoSuite also supports a growing user community through documentation and tutorial websites, workshops, Slack-based support, and the MyoChallenge competitions [24]–[27], which have provided community benchmark problems for neuromechanical control in manipulation, locomotion, and bionic human simulation.

MyoAssist was initiated following MyoChallenge 2024, “Physiological Dexterity and Agility in Bionic Humans” [26]. Motivated by user requests and community interest in the two assistive-device environments featured in the challenge, we packaged myoMPL and myoOSL as MyoAssist 0.1 [18], an initial MyoSuite sub-suite for musculoskeletal simulations with prosthetic and exoskeleton systems. The interest generated by this release motivated the development of MyoAssist 1.0 as a broader platform for physical human-device interaction.

### B. MyoAssist 1.0

MyoAssist 1.0 includes 15 assistive-device models and two training frameworks for controller development. A MyoAssist simulation environment assembles modular components into a human-device-task system (Section III). The training frameworks define how controllers are represented, trained, and evaluated in these environments (Section IV). As summarized in Fig. 1, the MyoSuite ecosystem provides the shared musculoskeletal modeling infrastructure, including models in the `myo_sim` repository; MyoAssist builds on this foundation through three repositories: `assist_sim`, which contains assistive-device models; `myoassist`, which contains the reinforcement-learning and controller-optimization frameworks, configuration files, examples, and utility code; and `myoassist.terrains`, a modular procedural terrain generator.

**Fig. 1.**
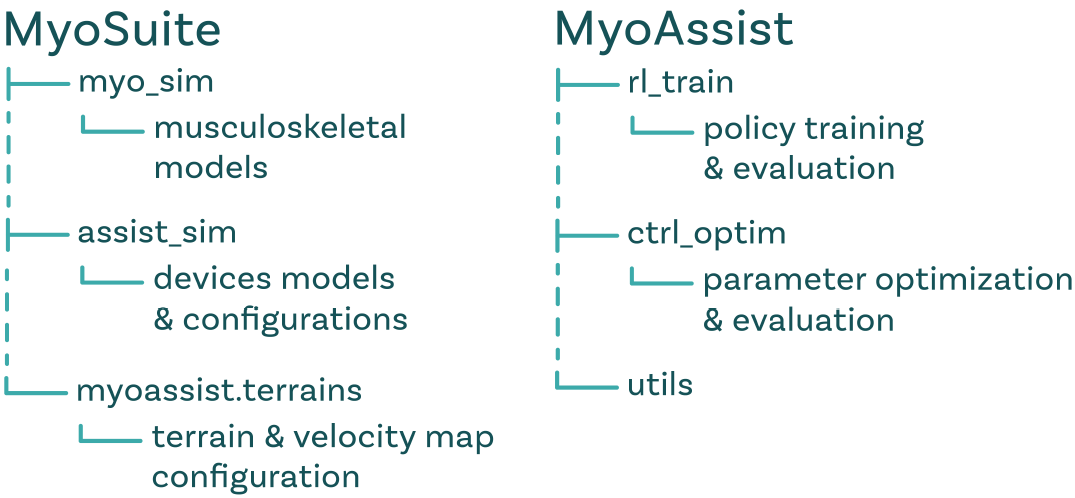
Software organization of MyoAssist 1.0 within the MyoSuite ecosystem.

This paper focuses on the platform organization, simulation environments, training workflows, and future outlook of MyoAssist; implementation details, hands-on tutorials, and source code are available through the following resources:

- MyoSuite myosuite website: https://sites.google.com/view/myosuite
- MyoSuite repository: https://github.com/MyoHub/myosuite
- MyoAssist tutorial website: https://myoassist.neumove.org/
- MyoAssist repositories:
  - https://github.com/neumovelab/assist_sim
  - https://github.com/neumovelab/myoassist
  - https://github.com/neumovelab/myoassist.terrains

A fixed snapshot of the software corresponding to MyoAssist 1.0 is archived as version 1.0.0 [1].

## III. Simulation Environments

MyoAssist simulation environments are organized as human-device-task systems that combine musculoskeletal models, assistive-device models, and task scenarios through a shared composition pipeline. Most assistive-device models are implemented as modular components that can be paired with compatible MyoSuite musculoskeletal models, although some environments require musculoskeletal-model modifications, such as segment removal for prosthetic limbs or specialized configurations for upper-limb and seated-mobility tasks. This section focuses primarily on gait-assistive devices because they constitute the majority of the current release and most clearly illustrate model-device compatibility, task variation, and environment construction.

### A. Musculoskeletal models

MyoAssist is designed to maintain compatibility with the musculoskeletal models available in MyoSuite. In this organization, MyoSuite provides biomechanically detailed human models, while MyoAssist extends these models with assistive devices and human-device simulation environments. Current MyoSuite models represent different body regions and modeling scopes, including the hand, arm, leg, torso, and full body. For gait-assistive simulations, MyoAssist supports models at multiple levels of complexity. Building on the 80-muscle *myoleg* model used in MyoAssist 0.1, the current release adds a 26-muscle lower-limb model and a 416-muscle full-body model [28]. These models allow users to choose the level of anatomical detail appropriate for their research question while balancing biological fidelity, interpretability, and computational cost.

### B. Assistive-device models

MyoAssist 1.0 provides simulation environments for real-world assistive devices (Fig. 2). The current release includes prosthetic and exoskeleton environments for gait assistance, upper-body environments for manipulation and trunk-support tasks, and a seated-mobility environment for wheelchair co-ordination. In most current exoskeleton environments, devices are modeled as rigid bodies with appropriate inertial properties, rigidly attached to musculoskeletal segments, with assistance applied directly at the corresponding biological joints. The University of Twente ankle-exoskeleton model instead uses compliant spring-damper attachments, such that device actuator torque is transferred to the musculoskeletal model through the resulting human-device interaction forces. MyoAssist also provides baseline device-controller templates for the lower-limb exoskeleton environments and the Open-Source Leg prosthesis, as described in Section IV-B. The simplified rigid-body, direct-torque representation provides a practical starting point for controller development and comparative simulation studies, while allowing more detailed human-device interface models, actuator dynamics, and device-specific controllers to be incorporated as the framework matures. For lightweight model inspection, the MyoAssist website also provides a browser-based interactive demo for exploring and interacting with a selection of device models, adapted from the MyoSuite demo and MuJoCo-WASM [29].

**Fig. 2.**
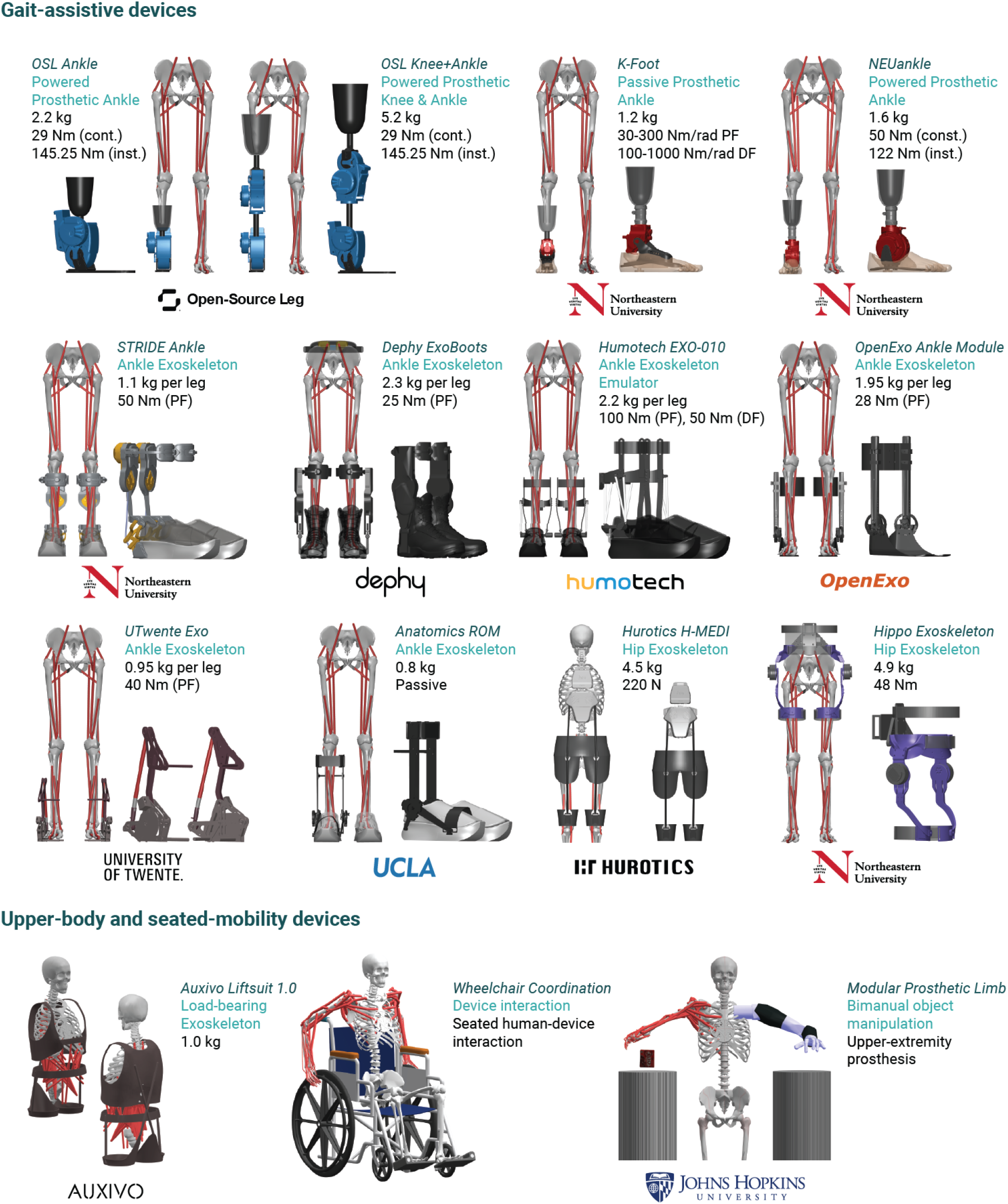
Device catalog of MyoAssist 1.0. The current release includes 15 assistive-device models spanning lower-limb gait assistance, upper-body manipulation and support, and seated mobility. PF and DF denote plantarflexion and dorsiflexion, respectively; cont. and inst. denote continuous and instantaneous torque ratings, respectively.

### C. Task scenarios

MyoAssist is compatible with the established MyoSuite task collection. Task scenarios refer to the non-device and non-musculoskeletal components that define what the human-device system is asked to do, such as terrain or scene geometry, target commands, episode initialization, and termination conditions. The MyoSuite task collection provides 13 task families, focused mostly on manipulation and locomotion, with variations in task details, difficulty levels, and physiological conditions, for a total of approximately 400 tasks that have been used across previous MyoSuite releases and MyoChallenge competitions. MyoAssist extends this collection with locomotion-specific scenarios for gait-assistive devices (Fig. 3), including configurable terrains composed of modular patches such as flat ground, slopes, stairs, and obstacles. These scenarios also support target-velocity maps that specify spatially varying desired walking speed and direction based on terrain layout and complexity, enabling studies of assistive control during speed- and direction-changing locomotion over configurable terrain.

**Fig. 3.**
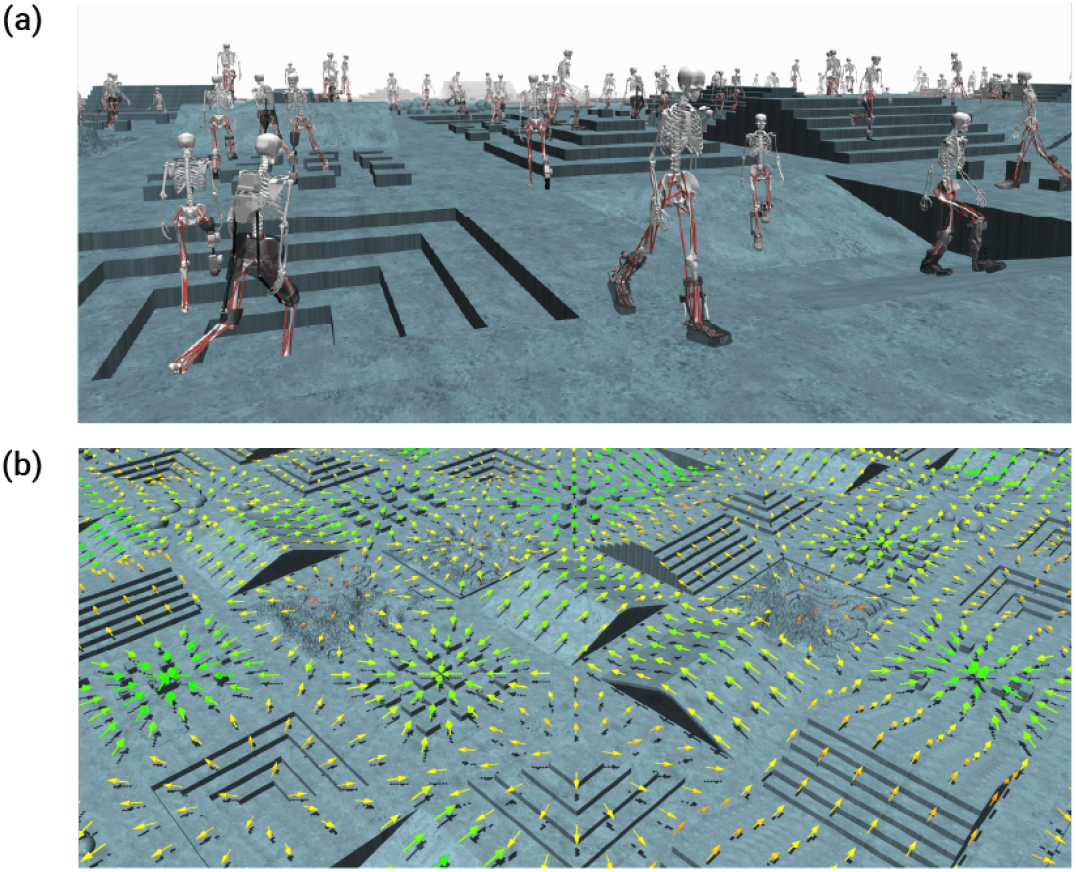
Locomotion-specific task scenarios in MyoAssist. (a) Configurable terrains are generated from modular patches. (b) Target-velocity maps define spatially varying desired walking speed and direction.

### D. Environment composition pipeline

Each lower-limb MyoAssist simulation environment is assembled from selected musculoskeletal, assistive-device, and task-scenario components (Fig. 4). This composition is implemented through MuJoCo’s `mjSpec` API, allowing MyoAssist to construct compiled MyoAssist simulation environments from modular components rather than requiring a separate hand-built model for each use case.

**Fig. 4.**
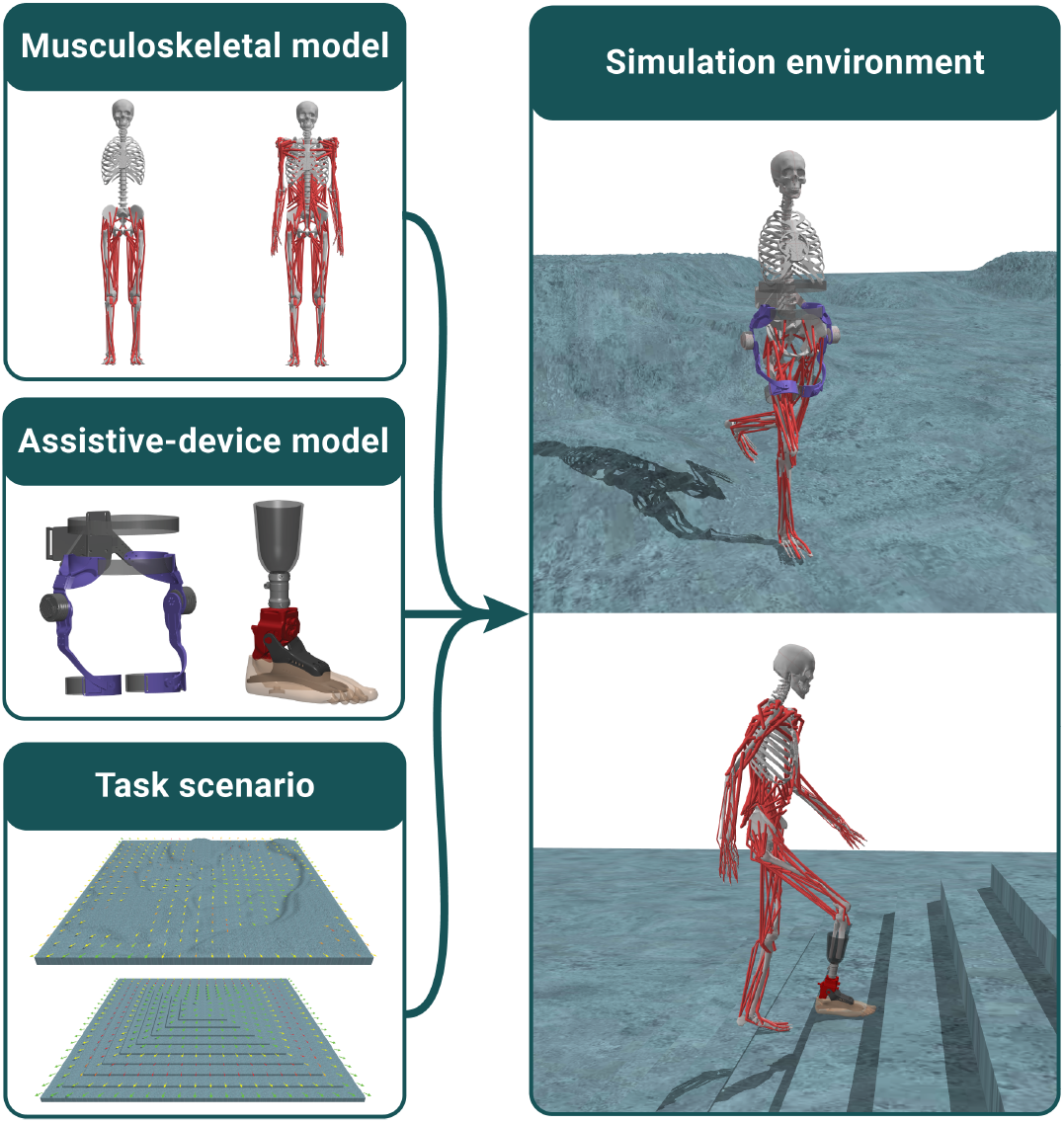
Environment composition pipeline in MyoAssist. Compatible musculoskeletal, assistive-device, and task-scenario components are combined to form a compiled human-device-task simulation environment.

Device integration is specified through a MuJoCo geometry XML together with a YAML configuration file. The YAML file defines the device attachment sites on the musculoskeletal model, any added joints or actuators, optional overrides to baseline model properties, and initial poses. The selected musculoskeletal, device, and task-scenario components are then merged into a single compiled MuJoCo model, which can also be exported to XML for inspection or reuse. This workflow supports both exoskeleton and prosthesis environments. In exoskeleton environments, device bodies are attached to an intact musculoskeletal model. In prosthesis environments, the relevant biological segments are removed or modified, residual-limb geometry is introduced, and the prosthetic device is attached in place of the removed anatomy. The resulting MyoAssist simulation environment provides a common interface for the reinforcement-learning and controller-optimization frameworks described in Section IV.

## IV. Training Frameworks

MyoAssist 1.0 provides two complementary frameworks for controller development in neuromechanical simulations of physical human-device interaction: a reinforcement-learning (RL) framework and a controller-optimization (CO) framework. Both frameworks operate on the same MyoAssist simulation environments, while framework-specific training configurations define how controllers are represented, trained, and evaluated. Broadly, the RL framework can be used to learn versatile and adaptive movement policies from reward-driven interaction, whereas the CO framework can be used to generate structured movement behaviors using interpretable control models. Together, these frameworks allow users to develop, compare, and extend learned and structured control strategies within the same MyoAssist simulation environments.

### A. Reinforcement learning (RL) framework

The RL framework provides a learned-control workflow for training policies in MyoAssist simulation environments (Fig. 5). In RL, a policy maps observations to actions at each control step and receives rewards and updated observations from the environment; policy parameters are then updated from batches of these interaction data (Fig. 5(a)). The framework connects this Gym-compatible interaction loop with configurable policy structures, task-specific training configurations, and standardized evaluation outputs, with implementation components organized around training entry points, environment wrappers, policy modules, reference data, evaluation tools, and saved results (Fig. 5(b)).

**Fig. 5.**
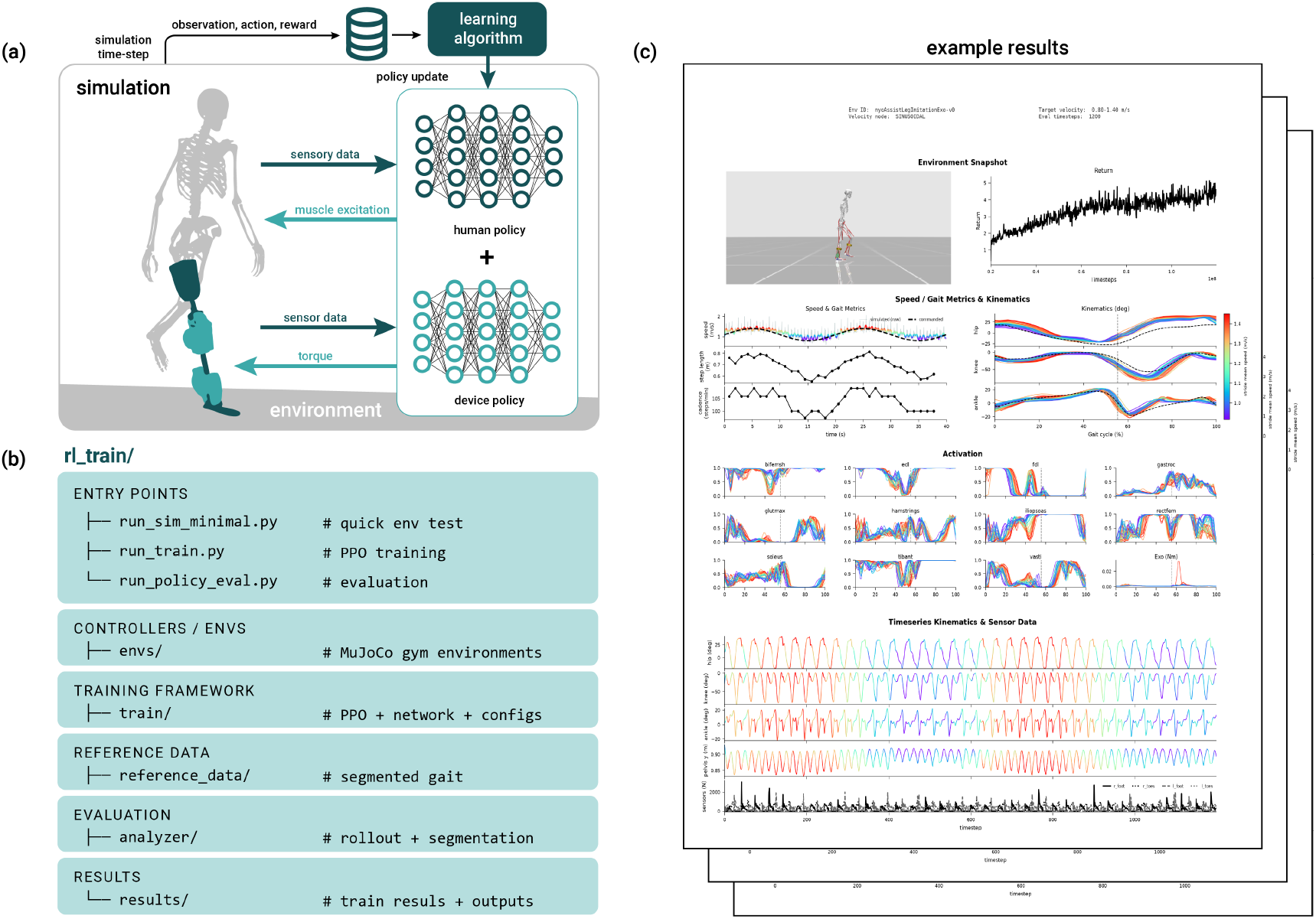
Reinforcement-learning framework in MyoAssist. (a) Gym-compatible RL loop, in which a policy maps observations to actions at each control step and is updated from batches of interaction data. (b) Organization of the RL training package, including training entry points, environment definitions and vectorization, reference data, evaluation tools, and saved results. (c) Representative evaluation outputs, including rendered snapshots, reward curves, gait kinematics and kinetics, muscle activations, sensor traces, and assistive-device outputs.

#### 1) Workflow

The RL framework follows the standard Gym interaction loop [23] used throughout MyoSuite, in which a MyoAssist simulation environment returns observations, a policy generates actions, and the environment advances the coupled musculoskeletal and assistive-device dynamics. The current implementation builds on the Stable-Baselines3 [30] implementation of proximal policy optimization (PPO) [31] and supports both single-environment execution for debugging and vectorized parallel execution for large-scale training. During training, the framework manages logging, checkpointing, periodic evaluation, and model export through the training configuration, allowing policy training runs to be inspected, resumed, and reproduced with minimal changes to the underlying simulation environment.

#### 2) Policy structure

The RL framework provides custom actor-critic policy structures for controlling muscle excitation, assistive-device actuation, or both. Policies can be implemented as a single actor that outputs the full action vector or as a compositional architecture in which separate human and assistive-device actors generate different action components while sharing a common critic. An indexing layer defines which observation entries are routed to each actor and how each actor output is mapped back to the full action space. This design allows the human and assistive-device controllers to use different observation sets and action mappings. For example, the human actor can be given proprioceptive and task-level observations, whereas the assistive-device actor can be restricted to measurements that approximate device-available sensing.

The framework also provides two optional left-right symmetry mechanisms for bilateral devices. First, left- and right-side assistive-device actors can share one set of weights while receiving equivalently ordered observations with the ipsilateral leg listed first, yielding mirrored device commands by construction [32]. Second, training can include a mirror-symmetry penalty that discourages the policy from producing inconsistent actions for mirrored states [33]. Either mechanism can be enabled independently through the training configuration.

#### 3) Training configuration

RL training configurations define the policy-learning problem, covering policy representation, controller-environment interfaces, task-scenario conditions, reward design, initialization, learning settings, and evaluation. They allow users to specify single or compositional policy architectures, actor-specific observation and action mappings, PPO hyperparameters, reference-motion data (when used), and previous-policy initialization. The reward function can be assembled from configurable terms such as velocity tracking, joint-position and joint-velocity imitation, end-effector imitation, muscle effort, activation smoothness, foot-contact loading, joint-limit forces, and step-level gait properties. For locomotion studies, target-velocity profiles can be uniform, sinusoidal, step-changing, or randomized across episodes, supporting training setups for speed-adaptive and speed-modulating policies. These configuration options allow users to implement motion imitation, reward-driven locomotion, transfer learning, and partial-observation studies without modifying the underlying MyoAssist simulation environment.

#### 4) Evaluation and reuse

The RL evaluation pipeline supports reproducible comparison and transfer of trained policies across MyoAssist simulation environments. Trained RL policies can be reloaded for evaluation, fine-tuning, transfer, or initialization of new training runs. The evaluation pipeline replays trained policies under user-defined conditions and generates standardized outputs for inspecting and comparing learned behaviors, including rendered videos, reward curves, gait kinematics and kinetics, muscle-activation traces, assistive-device output profiles, and saved gait data (Fig. 5(c)).

### B. Controller optimization (CO) framework

The CO framework provides a structured-control workflow that formulates controller tuning as parameter optimization in MyoAssist simulation environments (Fig. 6). As illustrated in Fig. 6(a), each candidate parameter set can define a human controller, an assistive-device controller, or both, and is evaluated in simulation to produce a scalar cost; Covariance Matrix Adaptation Evolution Strategy (CMA-ES) [34] then uses the resulting population of parameter-set–cost pairs to update its sampling distribution and generate new candidates for the next optimization generation. The framework connects this parameter-search loop with interpretable controller structures, configurable cost functions, and standardized evaluation outputs, with implementation components organized around optimization scripts, controller definitions, CMA-ES drivers, reference data, evaluation tools, and saved optimization results (Fig. 6(b)).

**Fig. 6.**
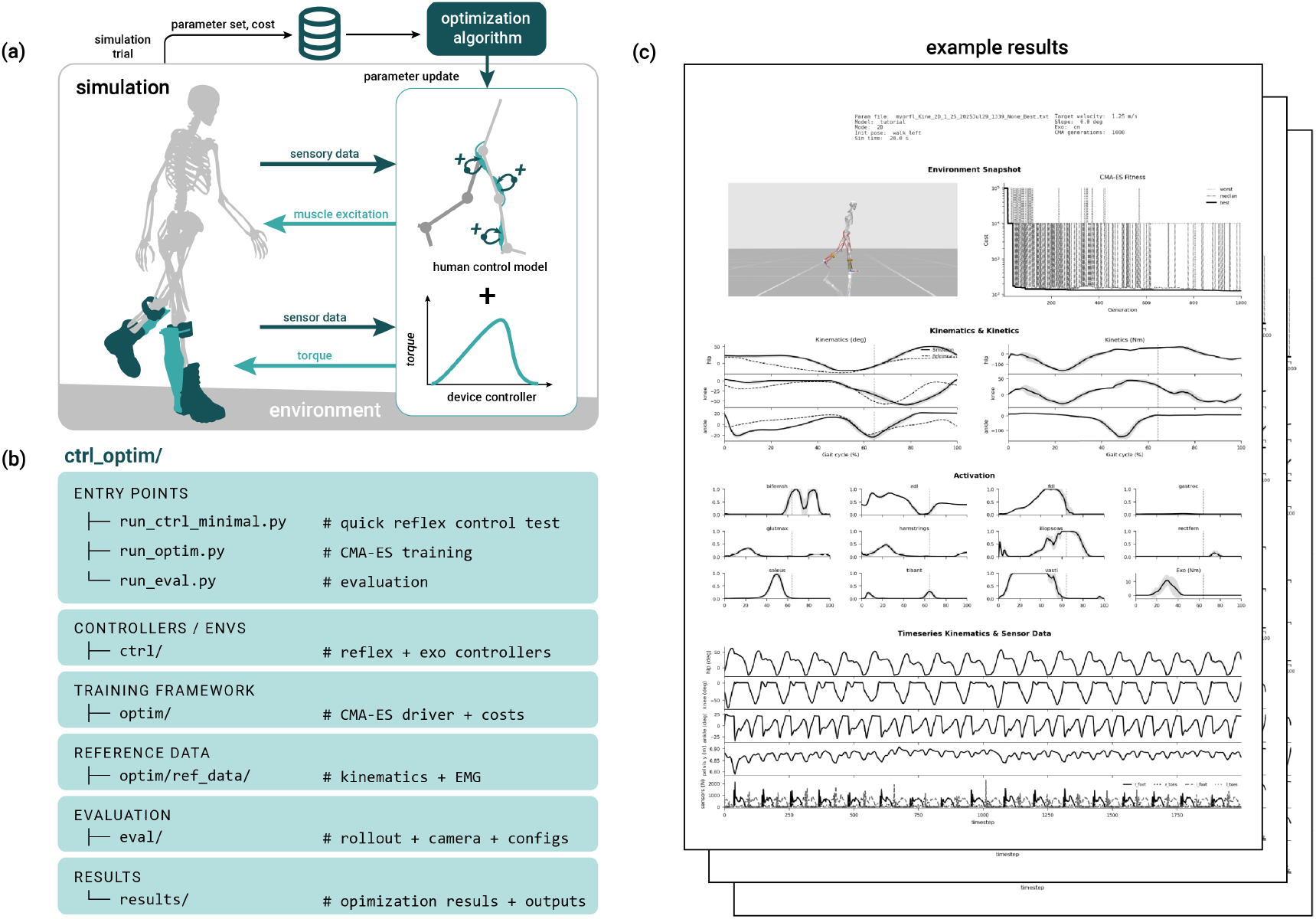
Controller-optimization framework in MyoAssist. (a) CMA-ES optimization loop, in which candidate human and/or assistive-device controller parameters are evaluated over simulation trials and used to update the sampling distribution. (b) Organization of the controller-optimization package, including controller definitions, CMA-ES drivers, reference data, evaluation tools, and saved optimization results. (c) Representative evaluation outputs, including rendered snapshots, optimization progress, gait kinematics and kinetics, muscle activations, sensor traces, and assistive-device torque profiles.

#### 1) Workflow

The CO framework tunes structured human and assistive-device controllers by simulating candidate parameter sets in MyoAssist simulation environments and assessing the resulting movement behavior with a configurable cost function. The current implementation uses CMA-ES, which samples candidate parameter sets, evaluates them in simulation, and updates the search distribution based on ranked outcomes. This workflow supports joint optimization of human and assistive-device controller parameters, optimization of either component alone, or evaluation of one fixed controller while tuning the other.

#### 2) Controller structure

The CO framework provides structured parametric controllers for both the musculoskeletal and assistive-device models. The human controller is a reflex-based neuromuscular locomotion controller based on Song and Geyer [35], with variants for three-dimensional and sagittal-plane locomotion control [36]. It generates muscle excitation from time-delayed sensory feedback, including muscle force, joint state, and ground-contact information. For exoskeleton simulations, MyoAssist provides a baseline exoskeleton controller that detects stance and swing phases and applies the widely used four-parameter spline-based torque assistance profile during the stance phase, with parameters defining peak torque and assistance timing [37]. MyoAssist also provides a generalized *N*-point spline controller, which extends this representation by allowing multiple torque magnitudes and timing locations and can be applied across assisted joints. For the powered prostheses, the framework provides a four-state impedance controller organized around early stance, late stance, early swing, and late swing [38].

#### 3) Optimization configuration

CO training configurations define the controller-optimization problem by specifying controller parameters, task-scenario conditions, optimizer settings, cost-function design, initialization, and evaluation. In the current framework, these settings include parameter bounds or initial values, CMA-ES hyperparameters, cost-function weights, and whether the human controller, assistive-device controller, or both are optimized. The baseline cost function uses a ranked, multi-stage structure suited to CMA-ES, a gradient-free, rank-based optimizer that can handle discontinuous simulation outcomes. The stages progressively distinguish early failure, constraint satisfaction, and final gait performance:

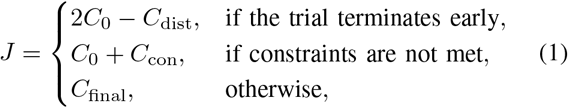

where *C*_0_ is a large constant that prioritizes earlier stages, *C*_dist_ rewards distance traveled before early termination, *C*_con_ penalizes constraint violations such as insufficient steadiness, speed mismatch, excessive ground-reaction forces, or joint-limit reliance, and *C*_final_ defines the final performance objective. This final objective can combine terms for muscle effort, kinematic tracking, velocity matching, ground-reaction-force limits, joint-limit reliance, EMG tracking, and gait symmetry. By modifying cost weights, feasibility criteria, or cost stages, or by introducing new cost terms, users can tune the same controller structure for different movement objectives, assistive-device configurations, and task scenarios.

#### 4) Evaluation and reuse

The CO evaluation pipeline supports reproducible inspection, continuation, and comparison of optimized structured controllers. Optimized controller parameters and optimizer states are saved so that CO runs can be inspected, resumed, or used to initialize new optimization runs. Best-parameter files can be reused across modified cost functions, task scenarios, assistive-device models, or musculoskeletal-model complexity levels, while saved CMA-ES states preserve the sampling distribution for continuation from an interrupted or partially completed run. The evaluation pipeline replays saved parameter sets in the corresponding MyoAssist simulation environment and generates standardized outputs for inspecting and comparing structured controllers, including rendered videos, optimization-progress plots, gait kinematics and kinetics, muscle-activation traces, assistive-device torque profiles, and cost breakdowns (Fig. 6(c)). The evaluation workflow is available through a JSON-configured command-line interface and is capable of batch processing.

## V. Discussion and Open-science vision

MyoAssist 1.0 establishes an extensible open-source framework for neuromechanical simulation of physical human-device interaction. The current release organizes each MyoAssist simulation environment as a composed human-device-task system consisting of compatible musculoskeletal, assistive-device, and task-scenario components, and provides reinforcement-learning and controller-optimization workflows for developing human and assistive-device controllers within a shared interface. This organization is intended to lower the barrier to constructing, comparing, and reusing simulation environments across prosthetic, exoskeleton, upper-body support, and seated-mobility applications, while preserving enough modularity for users to adapt models, task scenarios, training configurations, and evaluation workflows to their own research questions.

The musculoskeletal modeling and control capabilities available in MyoAssist will continue to expand as MyoSuite advances. MyoAssist is built around MyoSuite’s musculoskeletal models and task interfaces, and the framework can therefore benefit from continuing community development of the broader MyoSuite ecosystem, including expanded task scenarios, reusable policy libraries, and biologically plausible control models [36]. MyoAssist is now compatible with a 416-muscle full-body musculoskeletal model [28], while ongoing development of muscle models with compliant tendons will further expand the human-model foundations available for future MyoAssist simulation environments. Through this compatibility, new MyoSuite components can be incorporated into MyoAssist without requiring each assistive-device model or training workflow to be reimplemented from the ground up.

MyoAssist-specific development will focus on improving model fidelity and strengthening links to experimental data. First, future releases can support higher-fidelity physical human-device interaction models. The rigid human-device attachments and direct joint-torque representations used in most current models provide practical starting points for scalable controller development and comparative studies, while the University of Twente ankle-exoskeleton model provides an initial compliant-interface implementation. Future extensions may incorporate more detailed soft-tissue and attachment dynamics, actuator and sensor dynamics, and communication delays. Second, future development should strengthen links to experimental and patient data through validation examples, standardized benchmark datasets, reference motions, motion libraries, and assistive-device sensor and torque data, with the longer-term goal of supporting experimental or hardware-in-the-loop workflows for reproducible simulation-based research.

MyoAssist is intended to serve as an open-science platform where simulation components developed for individual assistive-device studies can be shared, reused, and extended. We invite research groups developing, modeling, controlling, or evaluating assistive devices to contribute new assistive-device models, task scenarios, controllers, and datasets through the shared interfaces of MyoAssist. By reducing unnecessary variation in the underlying musculoskeletal models, assistive-device model integration, and simulation infrastructure, this shared-platform approach can make assistive-device research not only more accessible, but also more reproducible and easier to validate.

## Acknowledgments

The authors thank Elizabeth Wilson and Elliott Rouse (Open-Source Leg); Kathryn Lee, Fatima Tourk, Nathan Carmichael, and Max Shepherd (Northeastern U); Matthew Mooney (Dephy); Carl Curran and Josh Caputo (Humotech); Riley Shepard and Zachary Forest Lerner (OpenExo); Michael Rose and Tyler Clites (UCLA); and Jaeha Yang and Giuk Lee (Hurotics) for their collaboration and technical support in incorporating the assistive-device models into MyoAssist, including sharing model resources, specifications, and clarifications. The authors also thank Olivia Cardillo and Jasmine Wang (McGill University), Roman Dowling and Emma Fleck (Northeastern University), and Herman van der Kooij (U Twente) for their contributions to the development and testing of MyoAssist.

## Notes

This work was supported in part by the National Institutes of Health (RF1AG096055).

https://myoassist.neumove.org

https://github.com/neumovelab/myoassist

### Competing Interest Statement

The authors have declared no competing interest.

